# THE DUAL ROLE OF THE *MICROCYSTIS AERUGINOSA* MICROBIOME ON CYANOTOXIN PRODUCTION: COMPETITION FOR AND REMINERALIZATION OF ORGANIC NITROGEN

**DOI:** 10.1101/2023.10.18.562967

**Authors:** Wei Li, David Baliu-Rodriguez, Sanduni H. Premathilaka, Sharmila I. Thenuwara, Jeffrey Kimbrel, Ty Samo, Christina Ramon, E. Anders Kiledal, Sara R. Rivera, Jenan Kharbush, Dragan Isailovic, Peter K. Weber, Gregory J. Dick, Xavier Mayali

## Abstract

Nutrient-induced blooms of the globally abundant freshwater toxic cyanobacterium *Microcystis* are the cause of worldwide public and ecosystem health concerns. The response of *Microcystis* growth and toxin production to new and recycled nitrogen (N) inputs, and the impact of heterotrophic bacteria in the *Microcystis* phycosphere on these processes are not well understood. Here, using microbiome transplant experiments, cyanotoxin analysis, and stable isotope tracing to measure N incorporation and exchange at single cell resolution, we monitored the growth, cyanotoxin production, and microbiome community structure of several *Microcystis* strains grown on amino acids and proteins as the sole N source. We demonstrate that 1) organic N availability shapes the microbiome community structure in the *Microcystis* phycosphere; 2) external organic N input leads to lower bacterial colonization of the phycosphere; 3) certain *Microcystis* strains can directly uptake amino acids, but with lower rates than heterotrophic bacteria; 4) biomass-specific microcystin production is not impacted by N source (i.e., nitrate, amino acids and protein) but rather by total N availability; and 5) some bacterial communities compete with *Microcystis* for organic N, but others remineralize organic N, in the process producing bio-available N for *Microcystis*. We conclude that organic N input can support *Microcystis* blooms and toxin production, and *Microcystis*-associated microbial communities play critical roles by influencing cyanobacterial succession through either decreasing (via competition) or increasing (via remineralization) N availability, especially under inorganic N scarcity.

## Introduction

Climate change (e.g., warming and changes in precipitation patterns) affects the frequency and severity of harmful algal blooms ^1^ (HABs) globally ^2^, including those of cyanotoxin-producing cyanobacteria that can contaminate drinking water ^3^. Lake Erie, the shallowest and warmest of the Great Lakes that are a vital freshwater resource, receives nutrients from urban, industrial, and agricultural sources. Since the mid-1990s, Lake Erie has experienced seasonal cyanobacterial blooms dominated by *Microcystis*, *Anabaena*, and *Planktothrix* ^4, 5^. *Microcystis,* one of the most globally abundant bloom-forming cyanobacteria, is recognized as the primary producer of cyanotoxins, including microcystins (MC) in Lake Erie ^6, 7^.

Cyanobacterial HABs (cyanoHABs) are usually linked to excessive phosphorus (P) and nitrogen (N) input ^8^. P overloading has long been widely recognized as a major contributor to phytoplankton biomass ^9, 10, 11^; however, N is emerging as a limiting nutrient in freshwater. This is especially the case during blooms, where the availability of N often restricts the growth of cyanobacteria ^12, 13, 14^. MCs are the major cyanotoxins produced by toxin-producing *Microcystis* and are N-rich compounds, containing about 14% N by mass, largely exceeding the average cellular N content in *Microcystis* dry mass (∼7%) ^15, 16^. It has been shown that N limitation favors non-toxic *Microcystis* strains and that N-limited *Microcystis* cells produce less MC, suggesting N plays a critical role in cyanotoxin production ^17, 18^. Unlike P, which is primarily found as phosphate, biologically available N occurs in a variety of different inorganic (e.g., nitrate, nitrite and ammonium) and organic forms (e.g., urea, amino acids [AAs], and proteins) ^19, 20^. Numerous studies have demonstrated the effect of adding inorganic N on *Microcystis* growth ^21, 22^, while dissolved organic N (DON) that can exceed 50% of the N pool in aquatic ecosystems ^23, 24^ is potentially an important N source for cyanobacteria^25, 26^. For example, in Lake Erie, the fraction of DON relative to the total N pool increases from early to late summer, with most of this DON likely produced autochthonously during the bloom. Phytoplankton, including *Microcystis,* rely on this DON later in the bloom, after depletion of inorganic nitrate early in the summer ^27^. Regenerated N in the form of ammonium has been shown to play an important role in the longevity of cyanoHABs in Lake Erie and other eutrophic lakes ^28, 29^. However, the mechanisms *Microcystis* might use to access this DON, which is a complex mixture of many organic molecules, remain mostly unknown.

Another manner that *Microcystis* might access organic N is through microbiome interactions ^30^, since *Microcystis* is a photoautotroph that provides fixed carbon to the microbial loop ^31^. Indeed, interactions between autotrophs and heterotrophs affect the growth of both organism types ^32, 33, 34, 35^. Recent studies have identified a unique bacterial community associated with *Microcystis* colonies, suggesting potential organic matter transfer between the cyanobacteria and heterotrophic bacteria ^22, 36, 37^. This *Microcystis-*associated microbiome contains highly-expressed metabolic pathways that could be used to break down DON, such as deamination or hydrolysis of peptide bonds that would result in liberation of ammonium ^36^. However, it is still unconfirmed if and how heterotrophic bacteria affect *Microcystis* N assimilation, particularly within the phycosphere of *Microcystis*, where significant phytoplankton-bacteria interactions take place ^31^. In general, despite the fact that N is a key nutrient affecting MC production in toxin-producing *Microcystis,* N dynamics in the phycosphere are poorly understood.

In the present study, we aimed to better understand the impact of the *Microcystis* microbiome on N acquisition and its subsequent influence on biomass accumulation and toxin production. We suspected that in *Microcystis*-associated bacterial communities, certain bacteria remineralize organic N that is subsequently incorporated by *Microcystis* for growth and/or toxin production, while some others may compete with *Microcystis* for N.

## Material and Methods

### *Microcystis* strains and microbiome transplants

We examined 6 cultures (Fig. 6B) derived from the following three *Microcystis* strains available from culture collections: *Microcystis aeruginosa* PCC7806 (PCC7806, “axenic”), and two Lake Erie isolates ^38^, *Microcystis* sp LE3 (“LE3”, xenic, single celled and toxic)^39^ and *Microcystis* sp LE19-8.1 (“LE19”; non-toxic and colonial) ^38^. These cultures included: 1) “axenic”: the axenic PCC7806, 2) “bacterized^+^”: PCC7806 added with the microbiome from toxic strain LE3, 3) “bacterized^-^”: PCC7806 added with the microbiome from nontoxic strain LE19, 4) “xenic”: the toxic LE3 strain with its native microbiome, 5) “transplant”: the toxic LE3 strain with its native microbiome to which we added the microbiome from the non-toxic LE19 strain, and 6) “hybrid”: the non-toxic LE19 strain with its native microbiome to which we added the toxic axenic PCC7806 strain. All batch cultures were maintained in BG-11 medium with 2mM nitrate in a growth chamber (Precision, ThermoScientific, CA) with constant temperature (22°C) and light (30 µmol photons m^-2^ s^-1^) under light:dark cycle (14 h : 10 h) ^40^ (referred as “standard cultivation condition” hereafter).

### Cyanobacterial growth with various N sources

Cultures were inoculated into N-free BG-11 or high N (2 mM nitrate), low N (50 µM nitrate), AAs (50 µM N) or protein (50 µM N); organic N concentrations were based on dissolved free AA concentrations measured in Western Lake Erie ^41^ and a study on a *Microcystis* strain isolated in Lake Taihu in China ^26^. Triplicate cultures were incubated in 12-well plates (Costar, USA) and cyanobacterial growth rates determined by *in vivo* chlorophyll-a fluorescence intensities measured every 2-3 days, using a plate reader (Cytation 5, Biotek, VT) (excitation = 440 nm; emission = 680 nm). Pair-wise cross-treatment comparisons of specific growth rates were carried out using Tukey’s Honest Significant Difference (Tukey’s HSD) method ^42^ (see Supplemental Material for more details).

### Microcystin quantification

Cultures grown on different N sources were collected at exponential growth phase for toxin analysis and particulate organic carbon measurement. Major congeners (i.e., MC-LR, D-Asp MC-LR and MC-HilR) were quantified using ultra-high performance liquid chromatography - mass spectrometry (UHPLC-MS) at University of Toledo following previously published protocols ^43, 44, 45^. To compare MC production between cultures, particulate organic carbon (POC) concentrations of each culture were determined using a TOC analyzer (Shimadzu TOC-L equipped with an SSM-5000 solid state module, Shimadzu, Japan) to normalize the MC concentrations. Detailed procedures can be found in the Supplemental Material. Tukey’s HSD method was carried out to compare the effect of microbiomes and N sources.

### Cell-specific incorporation of organic N

To quantify bacterial and cyanobacterial incorporation of organic N substrates, we carried out a nanometer-scale stable isotope probing (nanoSIP) approach ^46^. We spiked ^13^C and ^15^N labeled algal AAs or protein (Cambridge Isotope Laboratories, Inc., MA) into late-log phase, nitrate replete cyanobacterial batch cultures (final N from AAs and protein 164 and 222 μM, respectively) in triplicate and incubated under standard cultivation condition. Isotope imaging of subsamples collected at 24 and 48 h and a set of control samples (no-addition and killed) was performed with a CAMECA NanoSIMS 50 at Lawrence Livermore National Laboratory following the protocol described in previous study ^46^. The C and N isotopic ratios of cells identified in the isotope images were quantified using L’Image (http://limagesoftware.net) and were subsequently used to calculate C and N cellular incorporation from substrate (C_net_ % and N_net_ %; see Supplemental Material).

### DNA extraction, metagenomic sequencing and analyses

To assess the effect of N sources on microbial community structure in cyanobacterial cultures, genomic DNA was extracted from initial and exponential growth phase cultures grown on different N sources using a DNeasy PowerSoil Pro DNA isolation kit (QIAGEN, Germany) following the manufacturer’s instructions. DNA extracts were quantified via Qubit dsDNA quantification assays (ThermoFisher, MA), and amplicons of the V4 hypervariable region of the 16S rRNA gene, amplified with the prokaryotic universal primer sets encoding F515/R806 ^47^ were sequenced on a NovaSeq 6000 platform (Illumina, CA) at Novogene Co. Ltd. Shotgun metagenomic sequencing of the xenic and hybrid cultures was carried out on a NextSeq2000 platform (Illumina, CA) with 2×150 cycles at Lawrence Livermore National Laboratory. All raw reads were deposited as a NCBI BioProject (PRJNA931951).

The 16S rRNA gene amplicon sequencing data was processed using QIIME 2 (v2021.8) ^48^. Briefly, the feature table of amplicon sequence variants (ASVs) were generated using DADA2 pipeline ^49^ with taxonomic classification assigned based on SILVA database release 132 ^50^. Alpha diversity in the samples was assessed by observed ASVs and Chao1 index ^51^. The beta diversity was visualized via principal coordinate analysis (PCoA) based on weighted UniFrac distance metrics between samples. The shotgun metagenome sequencing analysis was carried out using a custom pipeline consisting of sequence qualification, assembly, binning, and taxonomic and functional annotation steps to generate and assess metagenome-assembled genomes (MAGs; see Supplemental Material).

### Enumeration of attached cells

Triplicate hybrid cultures with and without protein addition were stained live with Sybr Gold DNA dye (ThermoFisher, MA) at 2X final concentration for 20 min in the dark. Cultures were then carefully transferred into chambered coverslips (Ibidi GmbH, Gräfelfing, Germany) to preserve the integrity of the colonies. Suspended *Microcystis* colonies were imaged using a Nikon CSU-W1 SoRa spinning disk super-resolution microscope with a 100x objective (Fig. 1A representative images). Excitation and emission settings for Sybr Gold and chlorophyll autofluorescence were 488 nm, 525/36 nm and 640 nm, 705/52 nm, respectively. Heterotrophic bacteria associated with *Microcystis* were manually counted in the collected micrographs in ImageJ (version 2.3.0). The effect of N sources on attached bacteria was assessed using a Poisson regression modeling on the cell counts in R (version 4.0.2) ^52^.

## Results and Discussion

### Organic N alters the *Microcystis* microbiome and decreases phycosphere attachment

We first assessed the bacterial community changes in the *Microcystis* cultures when switched from nitrate to AAs or protein with 16S rRNA gene amplicon sequencing (4,566,558 reads after quality filtering, ∼3900 unique amplicon sequence variants, ASVs, from 26 prokaryotic phyla). We removed ASVs identified as *Microcystis* (range from 0.9% to 51.8% of total reads) in further analyses. The most abundant ASVs belonged to the phyla Proteobacteria, Bacteroidetes, Armatimonadetes and Actinobacteria, which together contributed over 90% of the total ASV counts in all samples after *Microcystis* ASV removal*. Pseudomonas, Rhizobium, Limnobacter, Sediminibacterium, Hydrogenophaga, Rhodobacteria, Sphingobacterium, Phreatobacter, Porphyrobacter, Gemmatimonas, Mariniradius, Curvibacter, Rosemonas, Methylophilus* and *Blastomonas* were the most abundant genera found at abundances >1 % in at least one sample. All these genera are commonly found in natural Lake Erie during blooms ^22, 38, 53, 54^, from which these cultures were derived. Taxonomic diversity varied across treatments, with Chao1 alpha diversity indices ranging from 42 to 774 species. The cultures growing on protein, except the xenic culture, hosted the least diverse microbial communities among the N sources tested. Particularly in the hybrid cultures, protein as the N source resulted in a significantly lower alpha diversity compared to nitrate as the N source (Tukey HSD, *p* <0.001). Similarly, in the bacterized^+^ and bacterized^-^ cultures, the alpha diversities of samples with protein as the N source were lower than those with nitrate and AAs (Tukey HSD, *p* <0.05).

Our analysis of community structure further indicates that bacterial composition in the cultures grown on AA were significantly different than those in the cultures grown on protein (R = 0.41, *p* = 0.0001, n = 30). Separation of microbial communities incubated with different N sources (Fig. 2) could be explained by the relative abundance changes of 6 genera, including *Sediminibacterium, Rhizobium, Limnobacter*, *Curvibacter*, *Pseudomonas* and a member of the Caulobacteraceae family. In both bacterized^+^ and bacterized^-^ cultures, samples with AA as the N source showed higher relative abundances of *Curvibacter* and *Pseudomonas*, while samples incubated with protein showed higher abundances of *Sediminibacterium*, *Rhizobium*, and *Caulobacteraceae* (Tukey HSD, *p* < 0.001). Furthermore, significantly higher relative abundances of *Sediminibacterium* (Tukey HSD, *p* = 0.032) and *Curvibacter* (Tukey HSD, *p* = 0.021) were observed in the AA treatment compared to the protein treatment in the hybrid cultures.

Our analysis of the hybrid and bacterized^-^ cultures enabled us to identify some changes in the microbiome likely driven by the presence of non-toxic colonies that appear to harbor unique taxa. The communities were both derived from the LE19 culture, but the hybrid also contained the LE19 *Microcystis* cells and their attached bacteria. We observed that bacterized^-^ and hybrid samples incubated with protein clustered closely (Fig. 2), whereas the AA-incubated samples were distinctly separated. We identified that mean relative abundances of ASVs closely related to *Sediminibacterium*, *Sphingobacterium* and *Roseomonas* in hybrid cultures grown on AA were 41.6, 10.7 and 5.7 times higher, respectively, than those in bacterized^-^ cultures grown on AA. We attribute these differences to the presence of the non-toxic *Microcystis* cells that likely support different bacterial populations, since they are non-toxic and form colonies (Fig. 1A) that provide a niche for biofilm attachment. We interpret the differential response of the microbiome to AA availability, unlike that for protein, to be caused by the increased availability of AA inside colonies due to the smaller size of AA compared to protein.

To examine the metabolic potential of the heterotrophic communities associated with *Microcystis* cultures, we generated a total of 22 high-quality MAGs including 2 *Microcystis* and 20 heterotrophic bacteria from the metagenomic libraries of the xenic and hybrid cultures grown on nitrate as the N source (Supplemental Table S2). We identified the gene homologs involved in major N metabolic pathways for each heterotrophic MAG and predicted the presence and absence of these pathways (Fig. 3). These communities grown on 2 mM nitrate as the N source shared many taxa, which was also revealed by 16S rRNA gene amplicon sequencing, and only a few MAGs generated in the xenic and hybrid metagenomic libraries were unique to one or the other (Fig. 3, Supplemental Table S2). Considering the N cycling metabolic capacity, the heterotrophic bacterial communities in both cultures possessed similar potential metabolic functions with a few exceptions. All MAGs have the capability to mineralize organic N to ammonium as well as synthesize glutamine, an essential AA, from ammonium. We also investigated glutamate synthesis pathways in the MAGs, as glutamate is a building block for MC molecules ^55^ and plays a critical role in a wide range of metabolic processes including carbon metabolism and the assimilation of ammonia into AAs ^56^. Roughly 50% of the MAGs are missing the complete pathway for glutamate synthesis from glutamine. These bacteria either synthesize glutamate with an uncharacterized pathway, or they require glutamate from other organisms. Furthermore, it has been shown that a natural phytohormone, indole-3-acetic acid (IAA), promotes *Microcystis* growth ^57^. We identified two MAGs containing the complete pathway of bacterial production of IAA given a source of tryptophan, suggesting these could be mutualistic bacteria that produce growth hormones to stimulate algal growth in exchange for fixed carbon from the algae ^32^. Regarding potential competition for inorganic N, one MAG was different than the others (LE19_11 from the hybrid culture) where we identified nitrate/nitrite transport and partial pathways involving dissimilatory nitrate reduction and nitrification. Generally, however, the metagenomic analysis did not indicate major differences in organic N cycling potential between the two microbial communities. Differences in the abilities of the two microbiomes to process and remineralize organic N are likely due to differential expression, or the presence of uncharacterized pathways. Direct measurements of N degradation activity and N incorporation over time are needed to confirm these communities exhibit distinct organic N remineralization rates.

Direct nitrate uptake by heterotrophic bacteria would require increased C incorporation from *Microcystis* compared to exogenous organic N and C uptake from AAs and protein, and transfer from autotrophs to heterotrophs has been shown to increase via direct attachment ^58, 59^. Thus, we generated the hypothesis that cultures growing on inorganic N would have higher levels of heterotrophic bacteria attachment to *Microcystis*. We tested this hypothesis with the hybrid culture in a subsequent experiment. We manually counted the number of attached heterotrophic bacteria on over 2500 *Microcystis* cells in micro colonies (<100 cyanobacterial cells per colony) via fluorescence microscopy sampled from the hybrid culture grown in media with nitrate only, nitrate + protein, and protein only (Supplemental Fig. S1). The Poisson regression models ^52^ generated from the bacterial counting data revealed that, in comparison to the nitrate only treatment, the number of bacteria attached to *Microcystis* cells decreased by 31% and 40% for nitrate + protein and protein only treatments, respectively (*p* < 0.001 (Supplemental Table S1). This suggests that when exogenous organic N is supplied, fewer heterotrophic bacteria rely on organic substrates produced by *Microcystis* and exuded into the phycosphere, likely at least partially explaining the detected decrease in attachment and shift of the bacterial community structure.

### Both N source and microbiome origin impact *Microcystis* growth

A few studies have suggested that certain *Microcystis* strains are able to uptake organic N compounds such as AAs ^26, 60^ and that heterotrophic bacteria can also remineralize organic N, enabling uptake of the released inorganic N by *Microcystis*^32^. However, to our knowledge the effect of the microbiome on utilization of organic N sources by *Microcystis* has not been thoroughly studied. Thus, we carried out growth experiments with combinations of different microbiome communities and *Microcystis* strains to demonstrate that both factors (i.e., the microbiome and *Microcystis* strains) combine to influence the ability of *Microcystis* to grow on organic N (Fig. 4). In our experiments, none of the three *Microcystis* strains grew in the N-free BG-11 medium, showing that our protocol to acclimate the cultures in N-free medium before the start of growth experiments successfully removed any intracellularly stored N. All PCC7806 cultures with or without added microbiomes exhibited detectable growth under all tested N sources, fastest with high nitrate (2 mM) and slowest with protein (Fig. 4), showing this strain can directly utilize organic N in both simple (AA) and complex (protein) form, and the growth rates in low nitrate and AAs were equivalent. The additions of microbiomes did not impact the growth rate of PCC7806 under nitrate or AAs (Tukey HSD, p *>* 0.05); however, with protein as the sole N source, the microbiome from LE19 (bacterized^-^), but not from LE3 (bacterized^+^), led to a significant increase in growth rate (Tukey HSD, p < 0.01). This shows that some, but not all, microbiomes can remineralize complex organic N for subsequent incorporation by *Microcystis*. Further, this suggests that bacterial community structure is a critical factor to determine the succession of *Microcystis* blooms if proteinaceous organic matter becomes the primary N source.

Unlike PCC7806 that exhibited modest growth responses to the introduction of a microbiome under different N sources, the LE3 culture exhibited stronger responses to the addition of another microbiome. The unaltered LE3 culture (xenic) only grew on high nitrate, with higher growth rate than PCC7806 (Tukey HSD, *p* < 0.001), but grew slower on low nitrate, and decreased in abundance with AAs or protein additions. However, upon addition of the LE19 microbiome (transplant), these cultures exhibited growth responses to organic N similar to the hybrid culture (Fig. 4A). This suggests a very different role of the two microbiome communities, with the community from the most recently isolated strain comprised of bacteria that remineralize organic N. Conversely, the LE3 strain isolated 25 years ago did not appear to include N remineralizers, and under protein as the only N source, these bacteria became antagonistic. Interestingly, shotgun metagenomic sequencing data revealed that both LE3 and LE19 microbiomes have a similar set of genes for known N metabolic pathways (Fig. 3). Since LE3 has been cultured in the laboratory, usually with excess nitrate (in BG-11 media), we suspect that the LE3 microbiome has lost the ability to regulate expression of genes to remineralize organic N, in contrast to LE19 that was isolated from Lake Erie relatively recently. Such interactions between algae and bacteria are common in aquatic environments ^31, 61^ and have been supported by a recent study on the *Microcystis* phycosphere in Lake Erie using metatranscriptomics showing that colony-associated heterotrophic bacteria expressed genes such as amino acid oxidases and deaminases as well as peptidases to regenerate inorganic N (i.e., ammonia) from organic N compounds ^36^.

### Microbiome-organic N interactions influence *Microcystis* toxin production

Since extracellular microcystin measured in the cultures were generally below the detection limit, we focused on intracellular toxin content. Microcystin-LR (MC-LR) was the predominant form of all MC congeners in all cultures, thus, we focused on this congener to examine treatment effects corresponding to the growth experiments discussed above. In the axenic cultures, the high nitrate treatment yielded significantly higher MC-LR production normalized to biomass compared to the other N sources (Tukey HSD, *p* < 0.05). MC-LR concentrations in low nitrate, AAs and protein treatments were indistinguishable from one another (Tukey HSD, *p* > 0.05) (Fig. 4B). This is consistent with previous studies showing that N limitation decreases MC-LR production ^18, 62^, but here we uncovered that this strain did not decrease its biomass-normalized MC-LR production when growing on organic N, even when growth was negatively affected (Fig. 4A and B). Transplanting the LE3 microbiome into this strain (bacterized^+^) led to an increase in MC-LR production under the low nitrate treatment, compared to both axenic and bacterized^+^ with AAs or protein as the N source. In contrast, the bacterized^-^ culture did not exhibit any statistical difference in MC-LR production among all tested N sources (Tukey HSD, *p* > 0.05). Further, MC-LR production in the bacterized^-^ culture under high nitrate was significantly lower than that in the axenic and bacterized^+^ cultures (Tukey HSD, *p* < 0.0001). Collectively, our experiments elucidated that in addition to N source, the microbial community associated with *Microcystis* influences the production of cyanotoxins. Specifically, the MC data suggest that the two microbiomes differentially impacted cell-specific toxin production under high and low N availability. One potential mechanism to explain this phenomenon is the impact of heterotrophic bacteria on the availability of different AAs for the *Microcystis* cells. First, heterotrophic bacteria from lakes can incorporate different AAs in a taxon-specific manner ^63^. Our metagenomic sequencing data showed several MAGs contain genes for AA transport, as well as N mineralization (Fig. 3), which may result in varying availability of AAs for other organisms. Second, several lines of evidence previously suggested that AA availability impacts cyanotoxin production. Tonk et al. suggested that availability and stoichiometric balance of leucine and arginine affected MC production in the cyanobacterium *Planktothrix agardhii* ^7^. Further, Dai et al. demonstrated that a *M. aeruginosa* strain preferentially incorporated certain AAs (e.g., alanine, leucine and arginine) over others, and its MC production was affected by the type of AA provided in the culture medium ^26^.

Another possibility to explain the effects of the interaction between the bacterial community and N sources on MC production is the hypothesized role of MCs in remediation of reactive oxygen species (ROS) stress ^64^. Assuming MCs play a role in ROS detoxification, the toxic LE3 (xenic) culture may not require its microbiome to detoxify ROS, whereas the non-toxic LE19 culture might. Thus, it is possible that the LE19 microbiome has a greater capability to remediate ROS stress than the LE3 microbiome does, although this is not apparent at the genome level. We found that many of the MAGs from heterotrophic bacteria in both LE3 and LE19 cultures included catalases and superoxide dismutase, showing that both microbiomes have the potential to detoxify ROS (Fig. 3). To compare ROS remediation across the two microbiomes, future experiments directly measuring catalase activities and expression of those genes will be needed.

### Competition for organic N in the *Microcystis* phycosphere

The data presented above suggest that bacterial community composition and the *Microcystis* strain type jointly influence the growth and toxicity response to alternative N sources such as AAs and protein. However, these data did not provide direct activity measurements of this response. To quantify direct N assimilation from organic N sources in the *Microcystis* phycosphere and distinguish the incorporation by *Microcystis* cells from that of associated heterotrophs, we incubated the *Microcystis* cultures with ^13^C and ^15^N labeled AAs and protein and traced the C and N isotope composition of individual *Microcystis* and heterotrophic bacterial cells using nanoSIMS (Fig. 5). We identified *Microcystis* and heterotrophic bacteria cells in the nanoSIMS images based on their distinctive morphologies (e.g., size and shape) (Fig. 5A). Across all treatments, we collected data from over 3000 *Microcystis* and 5000 heterotrophic cells after 0, 24 and 48 hrs of incubation, and calculated the percentages of cellular C (*C_net_%*) and N (*N_net_%*) biomass derived from the labeled substrates for those >8000 individual cells. Linear regression of median values of *N_net_%* vs. *C_net_%* (Fig. 5B-E) reveals that *Microcystis* cells incorporated 4.4 and 10.8 times more of its N quota compared to its C quota from AAs and protein, respectively, whereas the relative proportions in heterotrophic cells were 1.6 and 3.9, respectively. This shows that, unsurprisingly, the photosynthetic *Microcystis* cells, able to fix C, used AAs and proteins primarily for N, while the heterotrophs used them for both C and N. However, we also note that the incorporation of some C from the substrates suggests direct incorporation of organic N into *Microcystis* biomass rather than cross-feeding from another organism that might have remineralized the organic N into ammonium ^65^ .

The PCC7806 incubations (axenic, bacterized^+^ and bacterized^-^) enabled us to compare the net incorporation of C and N from AAs and proteins with and without heterotrophs. Heterotrophic cells in the bacterized^+^ and bacterized^-^ cultures were equally labeled (MWW test, *p* > 0.05) after 24 hrs (Fig. 5B), more so from N (*N_net_%* = 2.8 and 2.6 % for bacterized^+^ and bacterized^-^, respectively) than C (*C_net_%* = 1.8 and 1.5 %, respectively). However, it should be noted that C is roughly 5 times more abundant by mass in a cell; thus, there was still a greater flux of C into cellular biomass than N from these substrates. After 48 hours, the bacterial cells exhibited decreased isotope labeling compared to 24 hours, suggesting that in both cultures, the labeled AAs had been depleted after 24 hours and the heterotrophs were incorporating more unlabeled organic matter exuded from the *Microcystis* cells. We also found that the bacterized^-^ microbiome was less labeled than the bacterized^+^ microbiome (Fig. 5B and D), suggesting the bacterized^-^ microbiome may be more efficient at incorporating unlabeled *Microcystis* exudates that would dilute the isotope signal. The *Microcystis* cells from these incubations were also significantly isotopically labeled (Fig. 5C and E), though less than the heterotrophs (Fig. 5B and D). Unlike the heterotrophs that were less labeled after 48 hours compared to 24 hours, the *Microcystis* labeling increased between 24 and 48 hours, which suggests that even though the AAs had been depleted, enough C and N had been remineralized by the heterotrophs to support additional isotope incorporation by *Microcystis* cells. Unexpectedly, the axenic *Microcystis* cells were more labeled than the *Microcystis* cells in the presence of the microbiomes, and labeling also increased between 24 hours and 48 hours. This shows that PCC7806 could incorporate both C and N from AAs, which did not become depleted after 24 hours, but in the presence of heterotrophs, *Microcystis* was outcompeted by the heterotrophic bacteria. Results for N incorporation from protein were similar to the AA data, but C incorporation was not (it was detectable but comparatively low). This indicates that *Microcystis* PCC7806 directly incorporated N but little C from protein after extracellular degradation, and further N incorporation was a result of cross-feeding from heterotrophs that produced ammonium or AAs. As with the AA treatment, heterotrophs outcompeted *Microcystis* for protein N.

Since *Microcystis* LE3 did not exist as an axenic culture, it was not possible to determine if this strain, like PCC7806, could also directly incorporate AAs and protein. However, the data from incubations of the original xenic and the transplant cultures are consistent with our previous interpretation from the PCC7806 incubations that the LE19 microbiome is better adapted to incorporate *Microcystis*-derived exudates compared to the LE3 microbiome, and potentially also more efficient at incorporation of protein and AAs. First, the heterotrophs in the xenic culture were more isotope labeled than the heterotrophs in the transplant culture, both in the AA (Fig. 5B) and the protein (Fig. 5D) incubations. This suggests that the newly formed microbiome from LE19 was incorporating more unlabeled *Microcystis* exudates and thus diluting the isotope signal from the protein and AAs. Second, *Microcystis* cells were more labeled in the xenic culture compared to the transplant, again both in the AAs (Fig. 5C) and the protein (Fig. 5E) incubations, suggesting the LE19 microbiome in the transplant cultures was a better competitor for those substrates, leading to lower incorporation by *Microcystis*. The growth data (Fig. 3) indicate that LE3, which has been in culture for decades, is a faster-growing copiotroph compared to LE19. Therefore, our interpretation is that the LE3 microbiome in the xenic culture is not able to supplement the N requirement of the associated *Microcystis* cells via remineralization of organic N despite the presence of the required metabolic functions in their genomes (Fig. 2). Since *Microcystis* cultures were grown in full strength BG-11 media with 2mM nitrate in the stable isotope tracing experiment, high enrichment of ^15^N and relatively low enrichment of ^13^C in LE3 *Microcystis* cells supports the idea that heterotrophic bacteria provided remineralized N from AAs and protein for the autotrophic cells.

Our experimental transplant design also enabled us to test the impact of the presence of different *Microcystis* strains on the heterotrophic bacterial growth on organic N. For example, the isotope labeling of the heterotrophs in the hybrid culture was roughly an order of magnitude higher than in the bacterized^-^ culture. These two cultures both comprise the LE19 microbiome and *Microcystis* PCC7806, the only difference being that the hybrid also included the non-toxic and colonial LE19 *Microcystis* cells and their physically associated heterotrophs. These data suggest that having only the toxic PCC7806 *Microcystis* strain in the bacterized^-^ culture inhibited heterotrophic bacterial activity, presumably because this microbiome has not been maintained in culture with a toxic strain. On the other hand, in the hybrid culture including both the PCC7806 and the non-toxic colonial LE19, the bacteria became much more highly enriched from AAs and protein, suggesting the non-toxic *Microcystis* strain did not inhibit bacterial growth and subsequent N remineralization. It is unknown if MC-LR or other cyanotoxins were responsible for this inhibition of bacterial growth, which can be the topic of a future study.

### Implication for cyanoHABs in freshwater ecosystems

The results of our experiments using laboratory co-cultures of different *Microcystis* strains incubated with two different microbiomes under different N sources suggest that the microbiome may be just as critical as the *Microcystis* strain in determining the influence of organic N on cyanobacterial biomass and toxin production. Studies in Lake Erie and other locations suggest that nitrate has been fully drawn down during late-stage blooms and that microbial recycling of organic N sustains blooms ^13, 19, 27, 66^. Our finding that its microbiome can remineralize organic N is consistent with this hypothesis. In addition, heterotrophs appear to outcompete *Microcystis* for organic N, thus, we hypothesize that symbiotic relationships with heterotrophs may be necessary for the cyanobacteria to access the regenerated ammonium from organic N uptake by heterotrophs, while *Microcystis* provide organic C or other needed molecules to heterotrophs. Another potential mechanism is that *Microcystis* is known as a superior competitor for ammonium compared to other phytoplankton (e.g. Blomqvist et al., 1994; Glibert et al., 2016). Therefore, access to DON via their heterotrophic partners, as demonstrated here, could be primarily by conversion to ammonium (without direct exchange via mutualism), which would still allow the blooms to persist under very low ambient N concentration and the presence of competing phytoplankton. Regarding bloom toxicity, our finding that toxin production was not affected by N form but rather by total N availability agrees with a previous laboratory study that examined urea ^21^, although another study suggested organic N additions led to lower toxin production in the field ^68^. This points to the importance of limiting total N input, including organic N, to control toxicity in eutrophic lakes. The fraction of non-nitrate N in Maumee runoff into Lake Erie over the last few decades has increased, which has correlated with cyanobacterial biomass during blooms ^20^. Total N loading will likely affect *Microcystis* strain composition, phycosphere community composition, and toxicity, but the exact outcomes will likely depend on the timing of external N input and thus-far uncharacterized inter-organism interactions that result in seasonal community dynamics ^22^.

## Acknowledgements

This work was funded by Lawrence Livermore National Laboratory’s (LLNL) Laboratory Directed Research Development grant # 20-ERD-061 and performed at LLNL under contract DE-AC52-07NA27344. We thank M. Zavarin for use of the TOC analyzer.

---

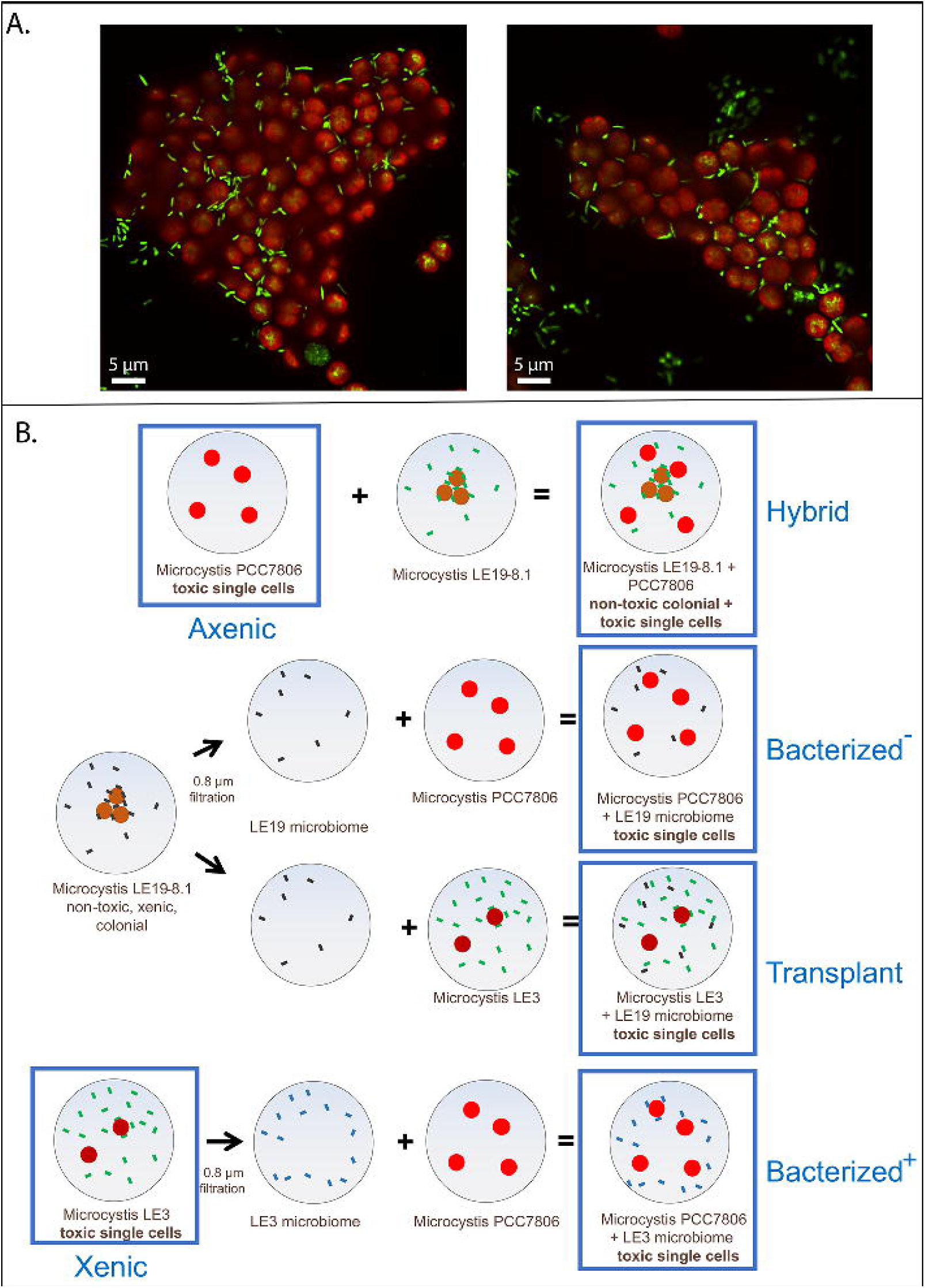

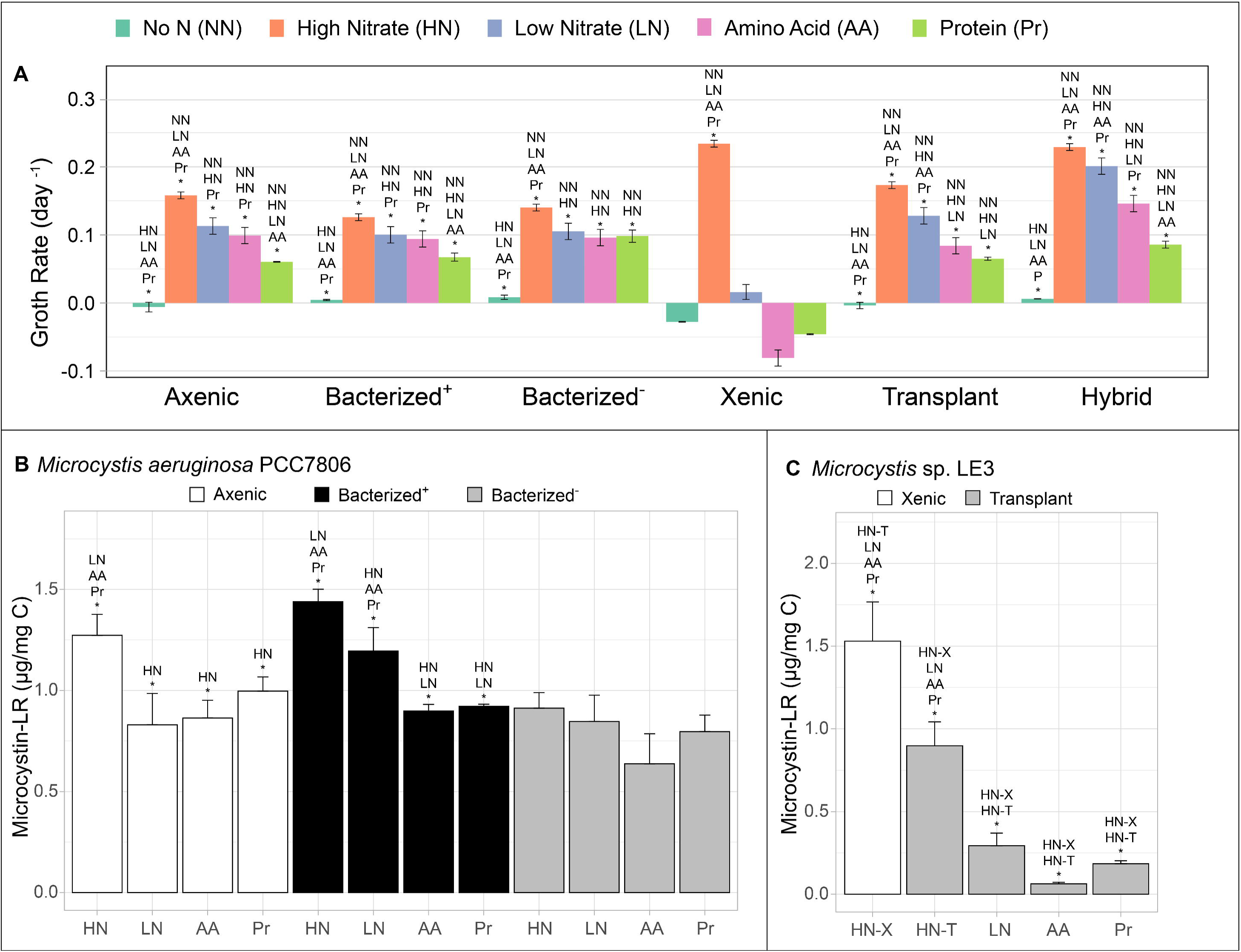

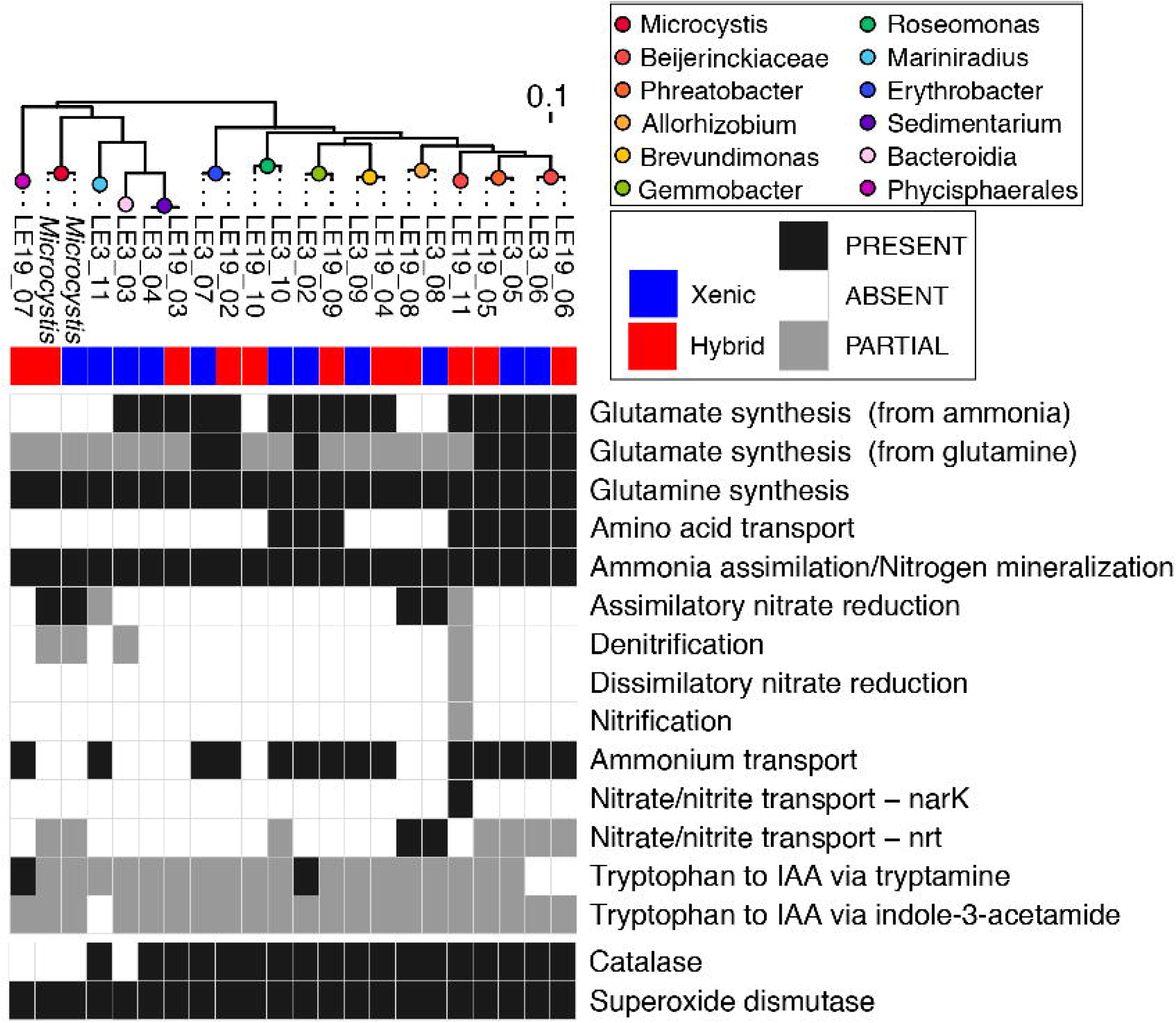

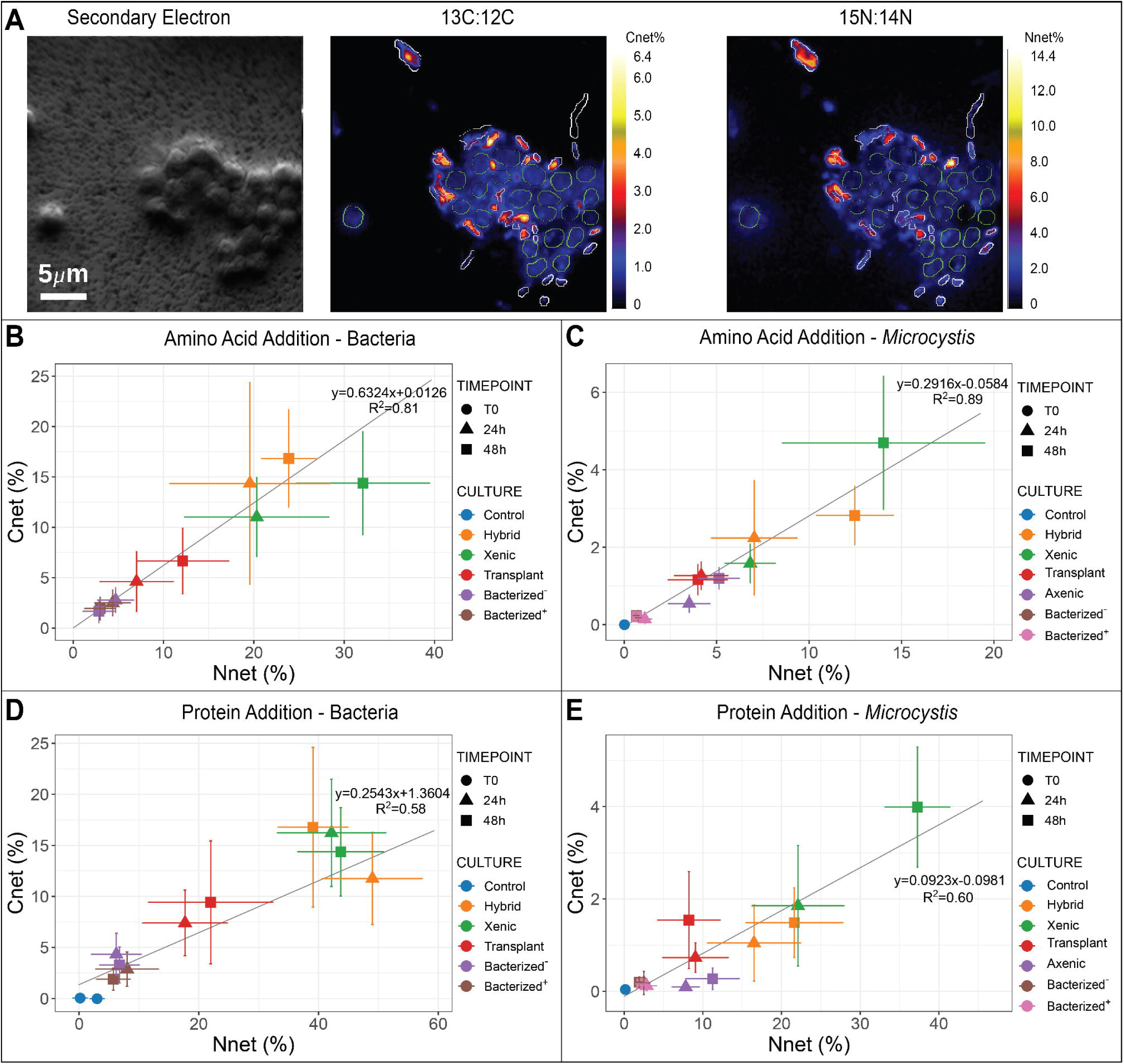

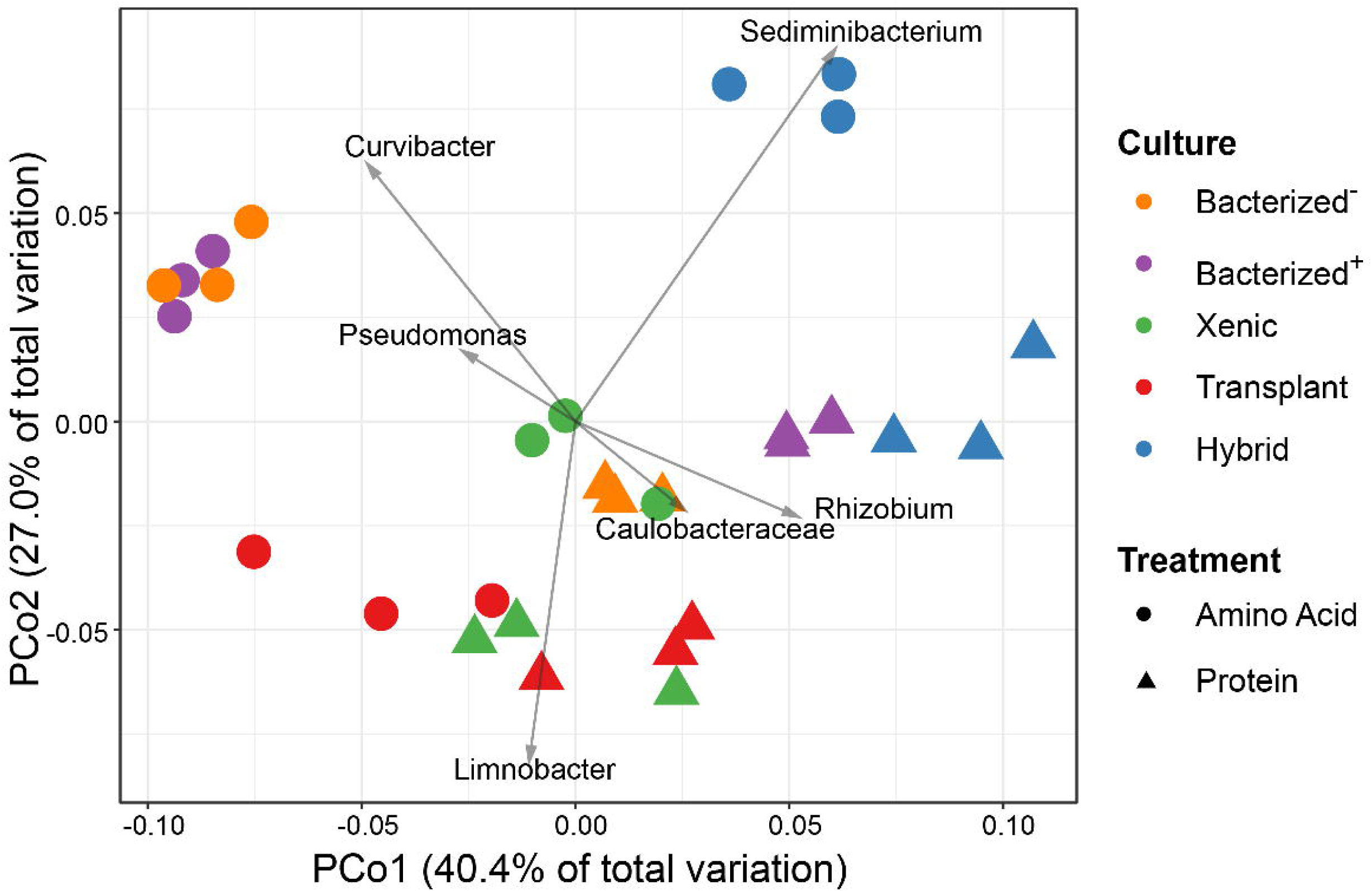

## Notes

### Competing Interest Statement

The authors have declared no competing interest.

