## Supplemental for "THE DUAL ROLE OF THE *MICROCYSTIS AERUGINOSA* MICROBIOME ON CYANOTOXIN PRODUCTION: COMPETITION FOR AND REMINERALIZATION OF ORGANIC NITROGEN"

**Supplemental Material**

Material and Methods

An optimized UHPLC-MS method was used for the analysis of MCs [1, 2, 3]. MCs were first separated using a Vanquish Flex ultra-high performance liquid chromatography (UHPLC) system (Thermo Scientific, San Jose, CA, US) equipped with a Waters HSS T3 C18 column (3.0 × 50 mm, 1.8 µm) and a guard column (2.1 × 5 mm, 1.8 µm). A binary gradient of $H_2O$ containing 0.1% HCOOH (mobile phase A) and $CH_3CN$ containing 0.1% HCOOH (mobile phase B) was used for chromatography. The flow rate was 0.667 µL/min and injection volume was 20 µL. The column compartment temperature was set to 45°C and column was equilibrated at 10% B. The gradient started with 10% B and was increased to 25% B in 0.03 minutes, to 46.4% in 0.97 minutes and to 95% in 1.70 minutes. After holding the gradient at 95% B for 0.50 minutes, the content of B was brought back to 10% in 0.12 minutes and maintained at 10% B for 2.68 minutes. MCs were identified using an Orbitrap Fusion Tribrid Mass spectrometer (Thermo) equipped with a heated electrospray ionization (ESI) source. Xcalibur (Thermo) software was used for data analysis. Samples were ionized in positive ion mode by heated ESI source set at 2400 V. Ion transfer tube temperature was 325ºC, and vaporizer temperature was 285ºC. Sheath gas was set at 40 arbitrary units (au) and auxiliary and sweep gas were set at 10 and 1 au, respectively. Selected ion monitoring (SIM)-MS with simultaneous tandem mass spectrometry (MS/MS) was used for screening and confirmation of MC ions. For quantitative analyses, all samples were analyzed by UHPLC-MS in triplicate and peak areas of extracted ion chromatogram (EIC) of monoisotopic MC ions were determined using Xcalibur. [1, 2, 3]

LC-MS-grade $H_2O$, $CH_3CN$, HCOOH and HPLC-grade $H_2O$, $CH_3CN$, HCOOH and $CH_3OH$ were purchased from Fisher Scientific (Pittsburgh, PA). Microcystin (MC) standards, MC-LR, D-Asp MC-LR, and MC-HilR, were from Enzo Life Sciences (Farmingdale, NY). Glass vials (20 mL) were purchased from DWK Life Sciences (Mainz, DE). Surfactant-free cellulose acetate (SFCA) membrane filters were from Fisher Scientific. Sep-Pak C18 cartridges were from Waters (Milford, MA). Glass vials (2 mL) and inserts (200 µL) were from Sigma (St. Louis, MO). The heated vacuum evaporator was from Eppendorf (Hamburg, DE).

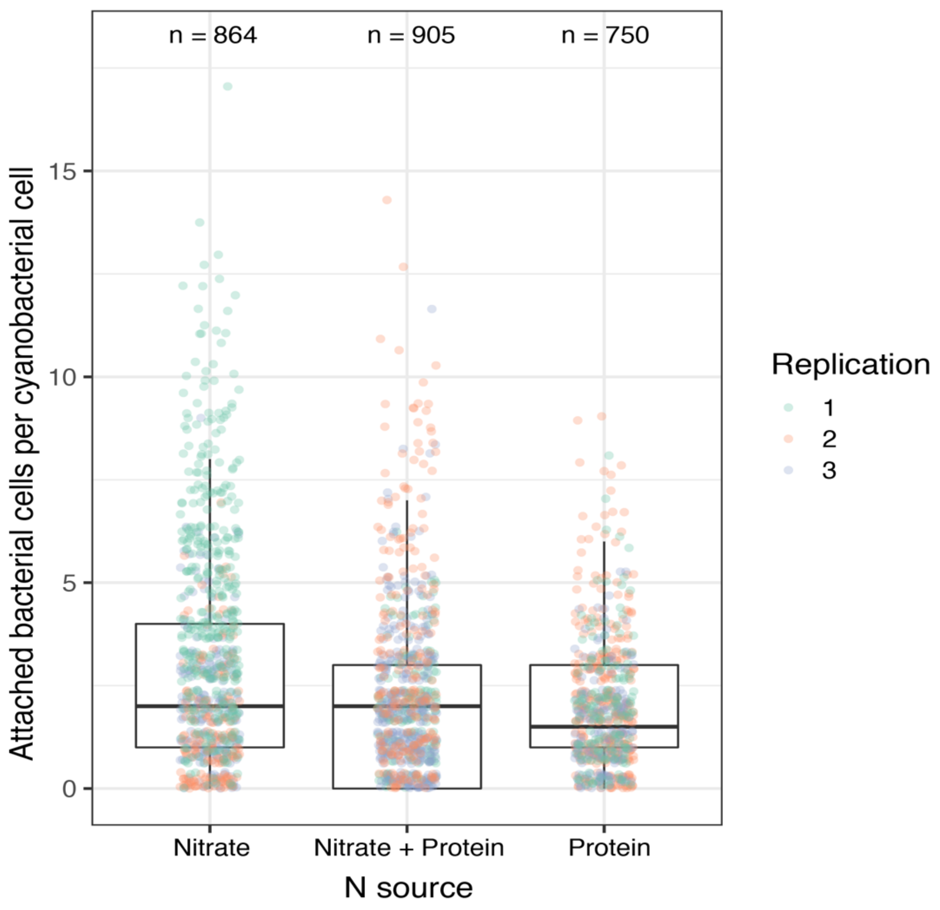


*Supplemental Figure S1. Attached heterotrophic bacteria counts in LE19 grown in nitrate, nitrate + protein or protein as N-sources.*

Supplemental Table S1. Poisson regression of attached bacterial counts of LE19 in different N sources. The mathematical form of Poisson regression model is $\log(y) = \alpha + \beta x$, where y is mean of counts in nitrate treatment (control), $\alpha$ is the intercept, x is the mean counts in nitrate + protein or protein treatments, $\exp(\beta)$ is the effect of nitrate + protein or protein treatments compared to nitrate (control).

| | Nitrate + Protein | | Protein | |
|---|---|---|---|---|
| | Estimate | *p* | Estimate | *p* |
| $\alpha$ | 1.0807 | $<2E^{-16}$ | 1.0807 | $<2E^{-16}$ |
| $\beta$ | -0.3662 | $<2E^{-16}$ | -0.5101 | $<2E^{-16}$ |
| $\exp(\beta)$ | 0.6934 | | 0.6004 | |

Supplemental Table S2. Summary of metagenomic assembled genomes (MAGs) generated from LE19-LLNL and LE3 culture shotgun metagenomic sequencing libraries.

| Cultrue | MAG | Completeness | Contamination | Contigs | Size (bp) | GC% | Contig L50 | Contig N50 | CDS | Cover M* | Lineage** |
|---|---|---|---|---|---|---|---|---|---|---|---|
| LE19-LLNL | LE19_01 | 99.7% | 0.3% | 447 | 4745655 | 42.5% | 52 | 27845 | 5279 | 78.5% | Bacteria;Cyanobacteria;Cyanobacteriia;Cyanobacteriales;Microcystaceae;Microcystis;Microcystis panniformis |
| LE19-LLNL | LE19_02 | 99.2% | 0.5% | 93 | 3393900 | 66.7% | 16 | 71798 | 3316 | 3.1% | Bacteria;Proteobacteria;Alphaproteobacteria;Sphingomonadales;Sphingomonadaceae;Erythrobacter |
| LE19-LLNL | LE19_03 | 99.0% | 0.0% | 18 | 2671600 | 36.1% | 2 | 398298 | 2543 | 1.7% | Bacteria;Bacteroidota;Bacteroidia;Chitinophagales;Chitinophagaceae;Sediminibacterium |
| LE19-LLNL | LE19_04 | 98.1% | 0.0% | 167 | 2646589 | 69.4% | 26 | 29149 | 2757 | 2.1% | Bacteria;Proteobacteria;Alphaproteobacteria;Caulobacterales;Caulobacteraceae;Brevundimonas |
| LE19-LLNL | LE19_05 | 96.0% | 0.3% | 107 | 3927339 | 67.7% | 19 | 57432 | 3913 | 1.4% | Bacteria;Proteobacteria;Alphaproteobacteria;Rhizobiales;Phreatobacteraceae;Phreatobacter |
| LE19-LLNL | LE19_06 | 95.4% | 0.9% | 89 | 4075456 | 66.3% | 13 | 93666 | 4001 | 1.6% | Bacteria;Proteobacteria;Alphaproteobacteria;Rhizobiales;Beijerinckiaceae |
| LE19-LLNL | LE19_07 | 94.3% | 0.0% | 78 | 3705307 | 67.2% | 16 | 79904 | 3186 | 4.3% | Bacteria;Planctomycetota;Phycisphaerae;Phycisphaerales;SM1A02;WM-009 |
| LE19-LLNL | LE19_08 | 94.2% | 2.2% | 592 | 5171102 | 61.2% | 111 | 14547 | 5660 | 0.9% | Bacteria;Proteobacteria;Alphaproteobacteria;Rhizobiales;Rhizobiaceae;Allorhizobium;Allorhizobium sp001713475 |
| LE19-LLNL | LE19_09 | 89.1% | 1.7% | 940 | 3906672 | 63.7% | 219 | 5515 | 4755 | 0.6% | Bacteria;Proteobacteria;Alphaproteobacteria;Rhodobacterales;Rhodobacteraceae;Gemmobacter |
| LE19-LLNL | LE19_10 | 70.6% | 1.5% | 1806 | 4208727 | 70.0% | 504 | 2698 | 5440 | 1.7% | Bacteria;Proteobacteria;Alphaproteobacteria;Acetobacterales;Acetobacteraceae;Roseomonas;Roseomonas stagni |
| LE19-LLNL | LE19_11 | 56.7% | 1.5% | 1584 | 3068889 | 67.1% | 498 | 2111 | 4337 | 0.5% | Bacteria;Proteobacteria;Alphaproteobacteria;Rhizobiales;Beijerinckiaceae |
| LE3 | LE3_01 | 99.9% | 0.1% | 242 | 4774342 | 42.8% | 45 | 32943 | 5242 | 67.5% | Bacteria;Cyanobacteria;Cyanobacteriia;Cyanobacteriales;Microcystaceae;Microcystis;Microcystis panniformis |
| LE3 | LE3_02 | 99.6% | 0.2% | 41 | 4103121 | 63.8% | 7 | 223502 | 4120 | 4.5% | Bacteria;Proteobacteria;Alphaproteobacteria;Rhodobacterales;Rhodobacteraceae;Gemmobacter |
| LE3 | LE3_03 | 99.0% | 0.5% | 21 | 3151917 | 36.4% | 3 | 249725 | 2740 | 1.5% | Bacteria;Bacteroidota;Bacteroidia;NS11-12g;UKL13-3;UBA6183 |
| LE3 | LE3_04 | 99.0% | 0.0% | 18 | 2691101 | 36.1% | 2 | 504466 | 2567 | 1.9% | Bacteria;Bacteroidota;Bacteroidia;Chitinophagales;Chitinophagaceae;Sediminibacterium |
| LE3 | LE3_05 | 98.6% | 0.0% | 161 | 4033918 | 67.6% | 24 | 43776 | 4074 | 2.1% | Bacteria;Proteobacteria;Alphaproteobacteria;Rhizobiales;Phreatobacteraceae;Phreatobacter |
| LE3 | LE3_06 | 98.3% | 0.9% | 131 | 4213910 | 66.3% | 18 | 65826 | 4180 | 3.4% | Bacteria;Proteobacteria;Alphaproteobacteria;Rhizobiales;Beijerinckiaceae |
| LE3 | LE3_07 | 97.4% | 0.4% | 173 | 3382380 | 66.6% | 33 | 34636 | 3387 | 2.0% | Bacteria;Proteobacteria;Alphaproteobacteria;Sphingomonadales;Sphingomonadaceae;Erythrobacter |
| LE3 | LE3_08 | 97.3% | 0.6% | 130 | 4977161 | 61.3% | 18 | 97796 | 5001 | 1.1% | Bacteria;Proteobacteria;Alphaproteobacteria;Rhizobiales;Rhizobiaceae;Allorhizobium;Allorhizobium sp001713475 |
| LE3 | LE3_09 | 95.8% | 0.2% | 259 | 2611814 | 69.2% | 44 | 17136 | 2818 | 3.0% | Bacteria;Proteobacteria;Alphaproteobacteria;Caulobacterales;Caulobacteraceae;Brevundimonas |
| LE3 | LE3_10 | 72.0% | 2.0% | 1338 | 4472359 | 69.6% | 346 | 4010 | 5286 | 0.7% | Bacteria;Proteobacteria;Alphaproteobacteria;Acetobacterales;Acetobacteraceae;Roseomonas |
| LE3 | LE3_11 | 56.5% | 2.8% | 1493 | 2556486 | 47.0% | 514 | 1782 | 3676 | 0.4% | Bacteria;Bacteroidota;Bacteroidia;Cytophagales;Cyclobacteriaceae;Mariniradius |

*CoverM, read coverage per-genome; **lineage based on GTDB database
